# The spatial and cellular portrait of Transposable Element expression during Gastric Cancer

**DOI:** 10.1101/2024.04.19.590342

**Authors:** Braulio Valdebenito-Maturana

## Abstract

Gastric Cancer (GC) is a lethal malignancy, with urgent need for the discovery of novel biomarkers for its early detection. I previously showed that Transposable Elements (TEs) become activated in early GC (EGC), suggesting a role in gene expression. Here, I follow-up on that evidence using single-cell data from gastritis to EGC, and show that TEs are expressed and follow the disease progression, with 2,430 of them being cell populations markers. Pseudotemporal trajectory modeling revealed 111 TEs associated with the origination of cancer cells. Analysis of spatial data from GC also confirms TE expression, with 204 TEs being spatially enriched. Finally, a network of TE-mediated gene regulation was modeled, indicating that ∼2,000 genes could be modulated by TEs, with ∼500 of them already implicated in cancer. These results suggest that TEs might play a functional role in GC progression, and highlights them as potential biomarker for its early detection.

## Introduction

Gastric Cancer (GC) is one of the leading causes of cancer death, with an estimated 800,000 demises per year ^1,2^. Although it has a global incidence, it is now considered an endemic malignancy in South America and some parts of Asia and Europe ^1,3^. The 5-year survival rate of advanced stages GC is ∼5%, while for early GC is >70% ^2,4^, highlighting the importance of detecting this malignancy in a timely manner ^3^. Persistent inflammation to the stomach is one of the main factors associated with the origin of GC. Particularly, chronic gastritis and Intestinal Metaplasia (IM) are precursor lesions, and the overall cascade progression of the disease can be recapitulated from Chronic Non-Atrophic Gastritis (NAG), Chronic Atrophic Gastritis (CAG), IM, early GC (EGC) and GC ^1,4–6^. Taking this into account, several groups have studied this progression using different modalities of RNA-Sequencing (RNA-Seq) to profile the changes in genome-wide gene expression ^6–8^.

RNA-Seq has remained the gold standard in gene expression studies due to its large-scale throughput, and nucleotide-level profiling of gene expression ^9,10^. Traditional RNA-Seq is also known as “bulk”, because it captures a homogenized portrait of gene expression ^11^. Although this has allowed for many advances in our knowledge, it has been somewhat limited for studying cancer due to intratumor heterogeneity ^12^. In turn, recent works using higher-resolution modalities of RNA-Seq, such as single-cell (scRNA-Seq) and spatially-resolved (srRNA-Seq), have been published ^6,13,14^. In addition to the identification of tumor subpopulations, scRNA-Seq allows for the reconstruction of cell trajectories through Trajectory Inference (TI) methods. TI corresponds to the modeling of dynamic cellular processes through the pseudotemporal arrangement of cells based on their transcriptional similarity ^15^. This methodology has been successfully studied to understand cancer evolution ^6,13,16,17^ and how different cancer subpopulations respond to treatment ^18^. In addition, by using this methodology, gene expression changes involved in cell fate decisions can also be studied, which can help understand what drives the changes in cell subpopulations towards a malignant genotype ^18–20^.

Transposable Elements (TEs) are genetic agents with the ability to move and increase their copy number ^21^. They are present in every eukaryotic genome known to date, and in humans they occupy about 50% of the genome ^22,23^. Broadly, they can be classified into retrotransposons, which transpose via an RNA intermediate, and into DNA transposons, which transpose via a DNA intermediate. Retrotransposons are further subdivided into Long Terminal Repeats (LTRs), Long Interspersed Nuclear Elements (LINEs) and Short Interspersed Nuclear Elements (SINEs), while DNA transposons are subdivided into DNA and Rolling-Circle (RC) TEs ^21,24^. Although most TEs are now genetically fixed, they can still become transcriptionally active, which can impact gene regulation ^25^. Current evidence supports their role as regulatory elements in health and disease ^26–28^. In particular, there are many examples of their global derepression and subsequent activation in a wide array of cancer types (reviewed in ^29^). Despite this, there are scarce works studying them using either single-cell or spatially-resolved methods. For example, only recently a group profiled gallbladder cancer using scRNA-Seq and applied TI to reveal that human endogenous retroviruses (HERVs, a type of LTR TEs) are associated with the transition of epithelial cells to the malignant status, and that they might act as regulators of gene expression in cancer cells ^13^.

Previously, I hinted at a regulatory role of TEs in EGC ^28^. An outstanding question in that study was whether TEs were expressed in previous timepoints, and whether their expression continues during GC. Here, by leveraging single-cell and spatially-resolved RNA-Seq data, I provide an extended analysis of the role of TEs in GC (**Fig. 1**). In this work, I show that TEs become up-regulated during the progression from gastritis to GC, and their expression is associated with the acquisition of malignancy. Additionally, network analysis suggests that they might be involved in the regulation of genes, highlighting a functional role in the GC cascade. Overall, these findings propose TEs as potential biomarkers for the detection of GC at its early stages.

**Fig. 1.**
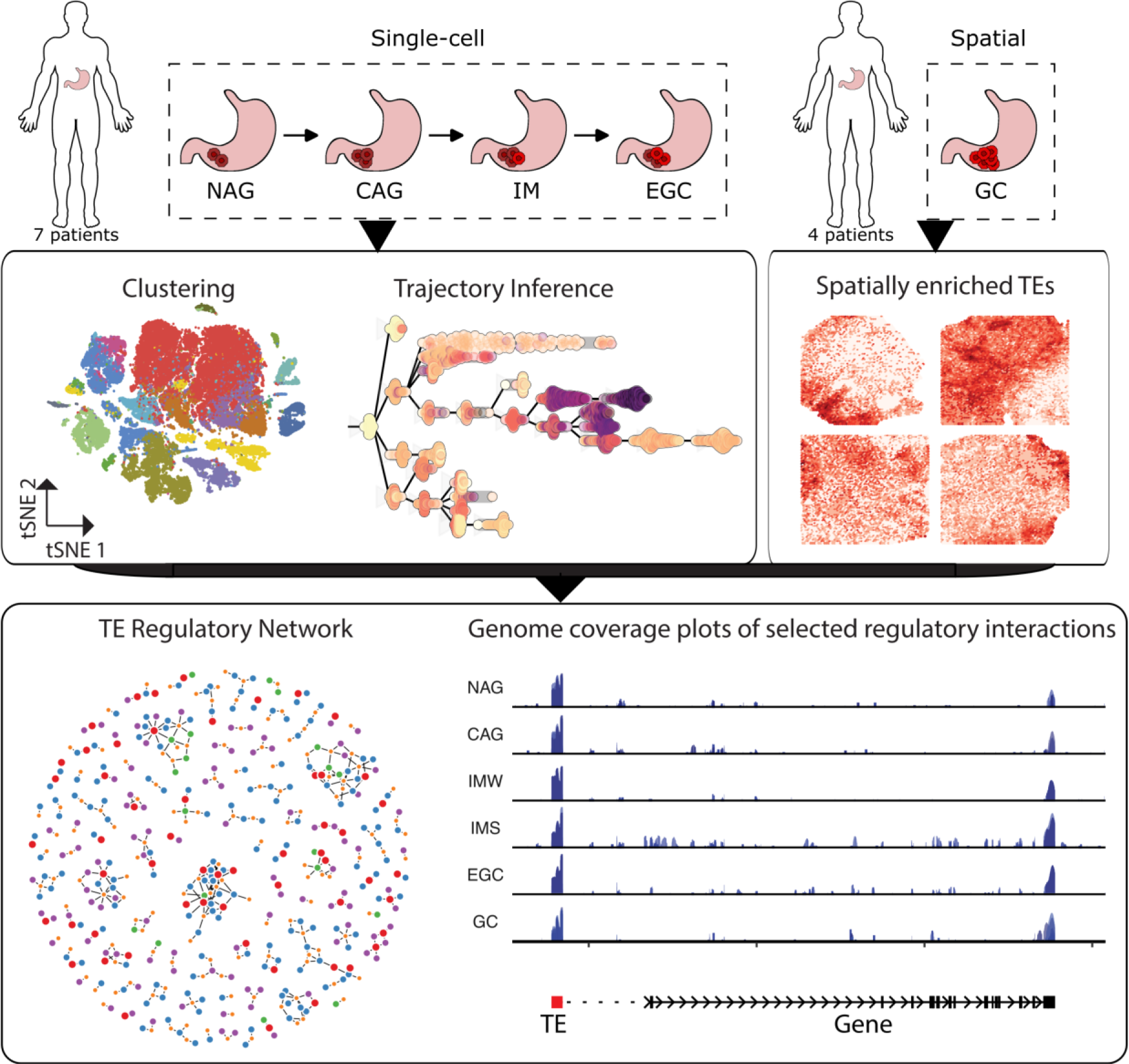
Overview of the analysis protocol used in this study. The single-cell RNA-Seq data used was obtained from 7 patients, and it spanned the progression from gastritis to cancer: Non-atrophic gastritis (NAG), Chronic atrophic gastritis (CAG), Intestinal metaplasia (IM) and Early gastric cancer (EGC). This data was first processed to obtain the dimensional reduction tSNE embedding, which was the basis for the identification and annotation of cell types, followed by a differential expression analysis to obtain the set of TEs characterizing the diverse cell populations. Then, Trajectory Inference methods were applied to predict the pseudotemporal progression of the cells along the EGC cascade. Additionally, Spatial RNA-Seq data from a different, independent study, obtained from 4 Gastric Cancer (GC) patients was used to further study TEs. Taking advantage of the pathologist annotations, TEs enriched in tumor regions were identified. Finally, the TEs identified from both analyses were used to build the TE regulatory network, which in turn allowed the prediction of genes whose regulation could be disrupted by TE expression.

## Results

### TE expression during the progression of gastritis to early GC

The single-cell raw sequencing data generated at the NAG (3 samples), CAG (3 samples), IM (3 samples, one wild and 2 severe) and EGC (1 sample) stages was aligned to the human genome. Afterwards, the resulting BAM alignment files were processed using SoloTE ^28^ to get matrices containing gene and TE expression per each cell. In order to get a preliminary overview on TE expression across the early GC cascade, a Principal Component Analysis (PCA) was carried out using a pseudobulk approach in which total expression was summarized at the sample level. This procedure was done 3 times, by using a matrix subsetted only to genes, another subsetted only to TEs, and one containing gene and TE expression (**Fig. 2a**). This analysis illustrates that gene expression alone recapitulates the differences between each time point, with the exception of the IM-Wild sample, which seems closer to the CAG samples.

**Fig. 2.**
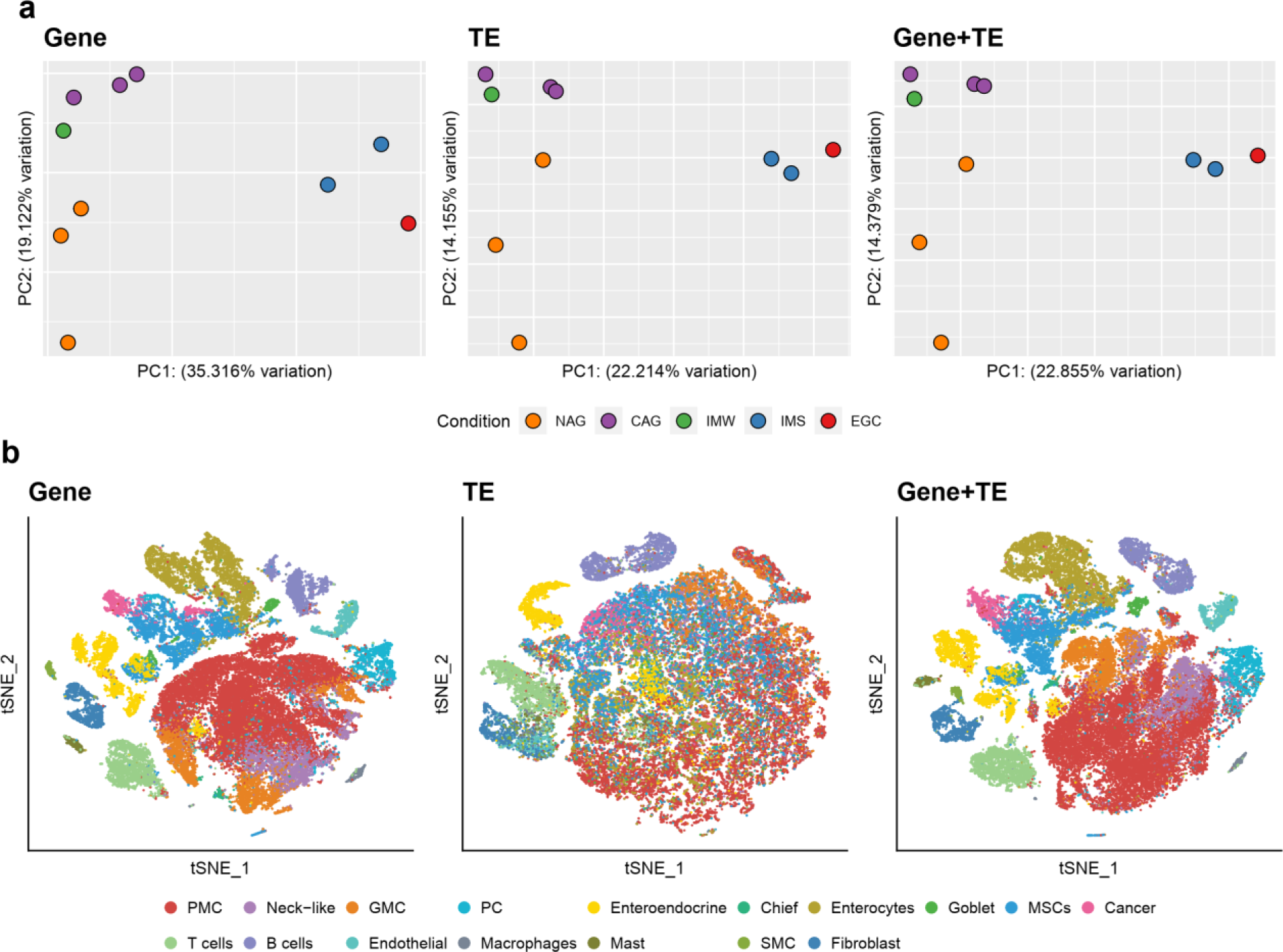
Dimensional reduction analyses of the progression from gastritis to early gastric cancer. **a.** Principal Component Analysis of the scRNA-Seq data, using gene expression only (first panel), TE expression only (second panel), and gene and TE expression (third panel). Each sample is color-coded according to the condition to which they belong. **b.** Dimensional reduction plots using the t-distributed stochastic neighbor embedding (tSNE). Similarly, the first panel shows the tSNE plot using gene expression only, the second using TE expression only, and the third using gene and TE expression. The plots are color-coded according to the cell types identified.

This suggests that only subtle changes occur during the IM-Wild timepoint. Using only TE expression, a similar distinction between timepoints can be seen. Although less variance is explained, the samples are still reasonably grouped in terms of their stage. As expected, the analysis using the complete Gene+TE matrix shows a result consistent with the ones done independently.

Next, to understand the influence of TEs at the cellular level, t-distributed stochastic neighbor embedding (tSNE) dimensional reduction analysis was carried out similarly, using the gene expression matrix, the TE expression matrix and the combined gene and TE expression matrix (**Fig. 2b**). The full Gene+TE matrix was processed first to generate a cell type annotation, resulting in 10 epithelial types, and 7 non-epithelial types. The epithelial population is comprised by Pit Mucous Cells (PMC), Neck-like, Gland Mucous Cells (GMC), Proliferative Cells (PC), Enteroendocrine, Chief, Enterocytes, Goblet, Metaplastic Stem-like Cells (MSCs) and Cancer cells. On the other hand, the non-epithelial population is comprised by T cells, B cells, Endothelial, Macrophages, Mast cells, Smooth Muscle Cells (SMC) and Fibroblasts. This cell type annotation was later transferred to the single-cell analyses done on genes and on TEs independently. In contrast to result obtained at the Gene level, the tSNE dimensional reduction of TE expression depicts more heterogeneity between cell types, in which PMCs, GMCs and MSCs seem to be more spread-out. It was previously shown that PMCs decrease and MSCs increase along the early GC cascade, and that there are transcriptional similarities between PMCs with both GMCs and MSCs ^6^. Thus, it can be suggested that TE expression could be indicative of intermediate cell states. Interestingly, the opposite can be seen in some non-epithelial subtypes, such as T cells and B cells. These cells exhibit a well-defined clustering pattern, hinting at a differential activation of a subset of TEs that might be modulating gene expression ^30^, with potential implications in T cell exhaustion ^31^.

Overall, these results show that although there is some variability in TE expression at the single-cell level, they are globally expressed throughout the progression from gastritis to EGC and recapitulate changes in each timepoint similar to those observed at the gene level.

### The single-cell expression of TEs in early GC

Given the global TE expression revealed by the previous analyses, I investigated what TEs could exhibit consistent or increasing expression from the premalignant gastritis status to EGC, and in what cell populations they might be enriched. To this end, the pseudobulked matrix was used, and a series of differential expression (DE) analyses were carried out using DESeq2^32^. Each of the CAG, IMW, IMS and EGC timepoints were compared to NAG, and TEs significantly up-regulated (having log2(Fold Change) > 0 and adjusted P-value ≤ 0.05) were selected. This resulted in a total of 2,581 TEs (Supplementary data 1). To also consider TEs that might already be activated in NAG, and whose expression remains constant throughout the EGC cascade (and thus, not appearing in the DE analysis), the TEs in the top 5% of highest expression across all stages were also selected (Supplementary data 1). This added 14,566 TEs to the set, resulting in a total of 17,147 TEs potentially associated with EGC progression (**Fig. 3a**).

**Fig. 3.**
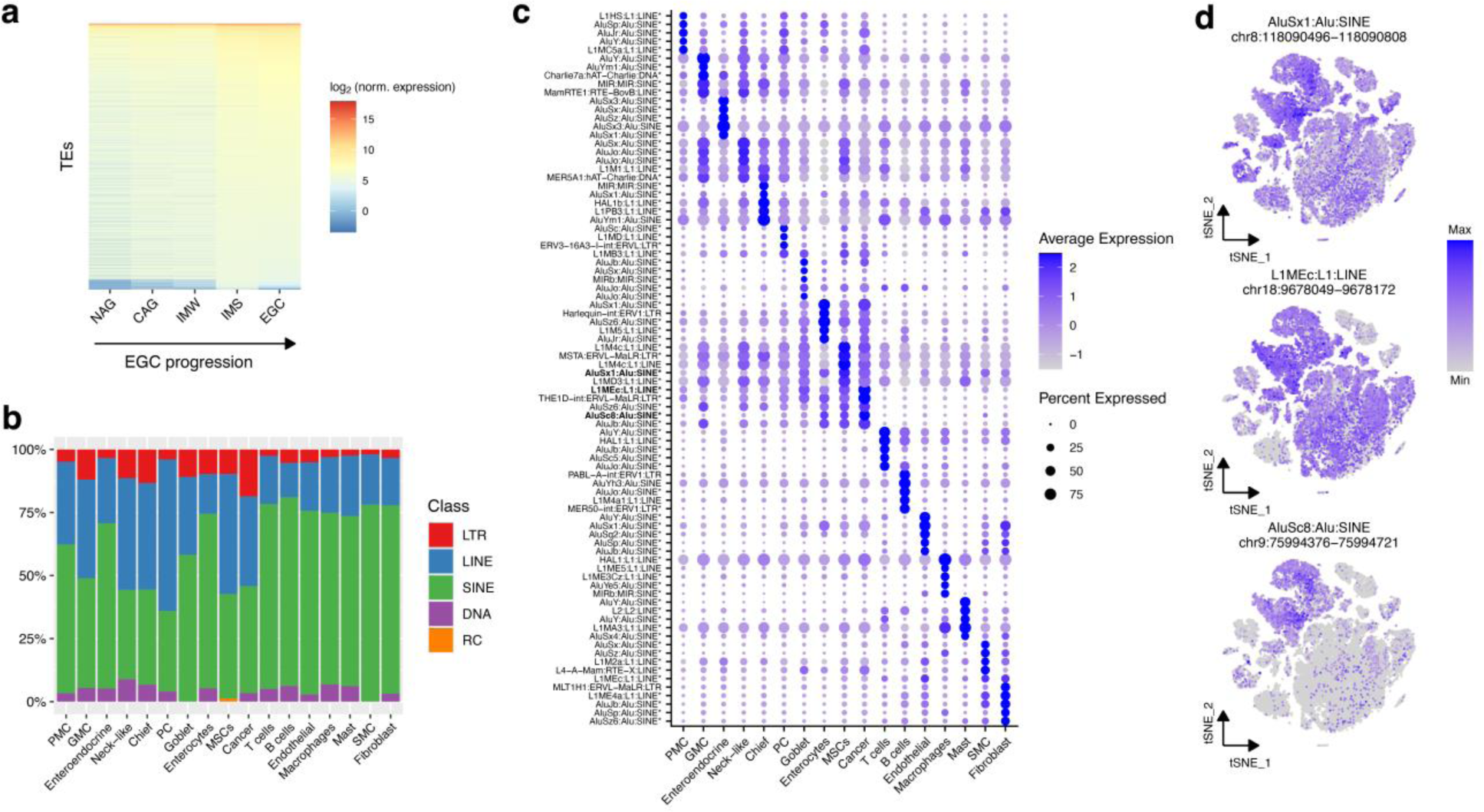
The single cell profile of TEs across EGC progression. **a.** Pseudobulked log_2_-normalized expression of TEs that were either differentially up-regulated at any time point with respect to NAG, or that were in the upper 5% at all time points. **b.** Class distribution of marker TEs of each cell type. **c.** Dot plot depicting the top 5 marker TEs of each cell type. In bold, TEs selected to show as example in the following tSNE plots. **d.** tSNE plots of selected TEs.

Marker analysis revealed that out of the TEs selected in the previous step, 2,430 have increased expression in the different cell populations (**Table 1**). Interestingly, a high proportion of these markers have locus resolution, which is essential to analyze their potential influence in gene regulation. Analysis of the major TE types revealed that they are all present to varying degrees in the set of markers (**Fig. 3b**), with a clear predominance of SINEs. Although previous works have pointed out a role of LINE L1 and LTR HERV TEs in cancer ^33^, there is evidence showing that SINE and DNA TEs are also involved by driving oncogene activation ^34^. This is also consistent with a recent work in colorectal cancer in which TEs were found to modulate gene expression and that SINEs were predominantly expressed ^35^.

**Table 1.** Marker TEs per cell cluster. For each cluster, the total number of marker TEs is shown, along with the number, and the proportion relative to the total, of locus-specific marker TEs.

| Cluster | Total marker TEs | Locus-specific marker TEs |
| --- | --- | --- |
| PMC | 61 | 55 (90.164%) |
| GMC | 92 | 73 (79.348%) |
| Enteroendocrine | 174 | 115 (66.092%) |
| Neck-like | 79 | 61 (77.215%) |
| Chief | 45 | 39 (86.667%) |
| PC | 25 | 22 (88.000%) |
| Goblet | 55 | 53 (96.364%) |
| Enterocytes | 133 | 126 (94.737%) |
| MSCs | 82 | 69 (84.146%) |
| Cancer | 59 | 53 (89.831%) |
| T cells | 356 | 312 (87.640%) |
| B cells | 227 | 209 (92.070%) |
| Endothelial | 214 | 158 (73.832%) |
| Macrophages | 163 | 135 (82.822%) |
| Mast | 163 | 158 (96.933%) |
| SMC | 146 | 129 (88.356%) |
| Fibroblast | 356 | 278 (78.090%) |
| **Total** | **2430** | **2045 (84.156%)** |

A caveat here is that the 10X single-cell data has a 3’ bias, and thus usually the terminal region of transcripts is captured and sequenced. Amongst TEs, the Alu family, part of the SINE group, has been reported to lead to truncated isoforms by acting as alternative transcription end sites ^36,37^. In turn, out of the 927 marker SINE TEs, 875 (94.4%) were part of the Alu family, suggesting that in EGC these TEs could be effectively acting as premature transcription end sites. It has been proposed that the impact of these events could be associated with disease progression ^36,37^, and indeed there is an example of such events in liver cancer, where Alu TEs were identified as the major TE becoming a terminal exon ^38^. Alternatively, the predominance of Alu TEs could be explained by their genetic structure: these elements harbor both a linker and a terminal A-stretch ^39^, which in turn could be causing internal poly-A priming during 10X 3’ single-cell sequencing.

Marker TEs, which are defined as TEs with high expression and expressed in a high proportion of the cells of a specific type, could be classified in 2 groups based on their percentage expressed in the remaining cell types (**Fig. 3c**, Supplementary data 2): group 1 – high percentage expressed; group 2 – low percentage expressed. For example, some of the top PMC, MSCs, Cancer, Chief and Neck-like markers belong to group 1, while the top T cells and B cells markers belong to group 2. In turn, group 1 would define markers that have a gradient of expression, and this pattern seen in MSCs and Cancer cells would be in agreement with the evidence suggesting that MSCs might give origin to GC. Similarly, neck cells differentiate into chief cells, and marker expression seem to coincide with this: the top neck cell markers are broadly expressed, while the top chief cells markers seem a bit more specific. To further illustrate this, tSNE dimensional reduction expression is depicted for 3 examples (**Fig. 3d**): AluSx1, located in chr8:118090496-118090808; L1MEc, located in chr18:9678049-9678172; and AluSc8 located in chr9:75994376-75994721. AluSx1 can be classified to group 1 considering that it is expressed in a high number of cells, but enriched in MSCs. The same argument applies to L1MEc with high expression in Cancer cells, but expressed at lower levels in other cells. Finally, AluSc8, also appearing as a Cancer marker, shows expression restricted to the region comprised by Cancer, MSCs and Enterocytes. It is worth noting that the expression of these TEs was measured with locus resolution (i.e., having only uniquely mapped reads), making unlikely that the observed expression heterogeneity could be attributed to ambiguity in read assignment. As suggested earlier, such gradient of TE expression spanning from PMC, GMC to MSC and then Cancer could suggest intermediate status between these cell types that could be mediated by TEs. It has been reported that as EGC progresses, PMCs decrease and MSCs increase, and that they have some transcriptional similarity ^6^, though it is unclear whether some PMC cells might be undergoing a malignant transformation to MSCs. If this is the case, these results would suggest that TEs are playing a role in that event.

Altogether, the evidence presented shows that TEs are expressed throughout the progression from gastritis to GC, and that their expression characterizes the different cell types, potentially highlighting intermediate status. In addition, the high proportion of TEs identified with locus resolution suggest that TEs becoming transcriptionally active in EGC have accumulated discriminative mutations, allowing the unambiguous assignment of sequencing reads. In turn, this would support the idea that TEs are playing a role in EGC progression via epigenetic polymorphisms, where changes in the transcriptional activity of fixed TE copies characterize cellular differences ^33^.

### TE expression is associated with the origin of cancer cells

After getting a global overview of TE expression, and assessing their cellular profile, I then asked if their expression is associated with the origin of cancer cells. To this end, I applied Trajectory Inference (TI) to model the pseudotemporal progression of cells from the normal to the malignant status using the *dynverse* R package ^15^. Briefly, this package evaluates more than 50 TI methods and identifies the one most suitable to the dataset, which in this case was PAGA-Tree ^40^. Then, *dynverse* represents the trajectory topology in a network of milestones where the cells are placed. The resulting trajectory was rooted at the milestone containing the highest number of NAG cells, and further analyzed (**Fig. 4a**).

**Fig. 4.**
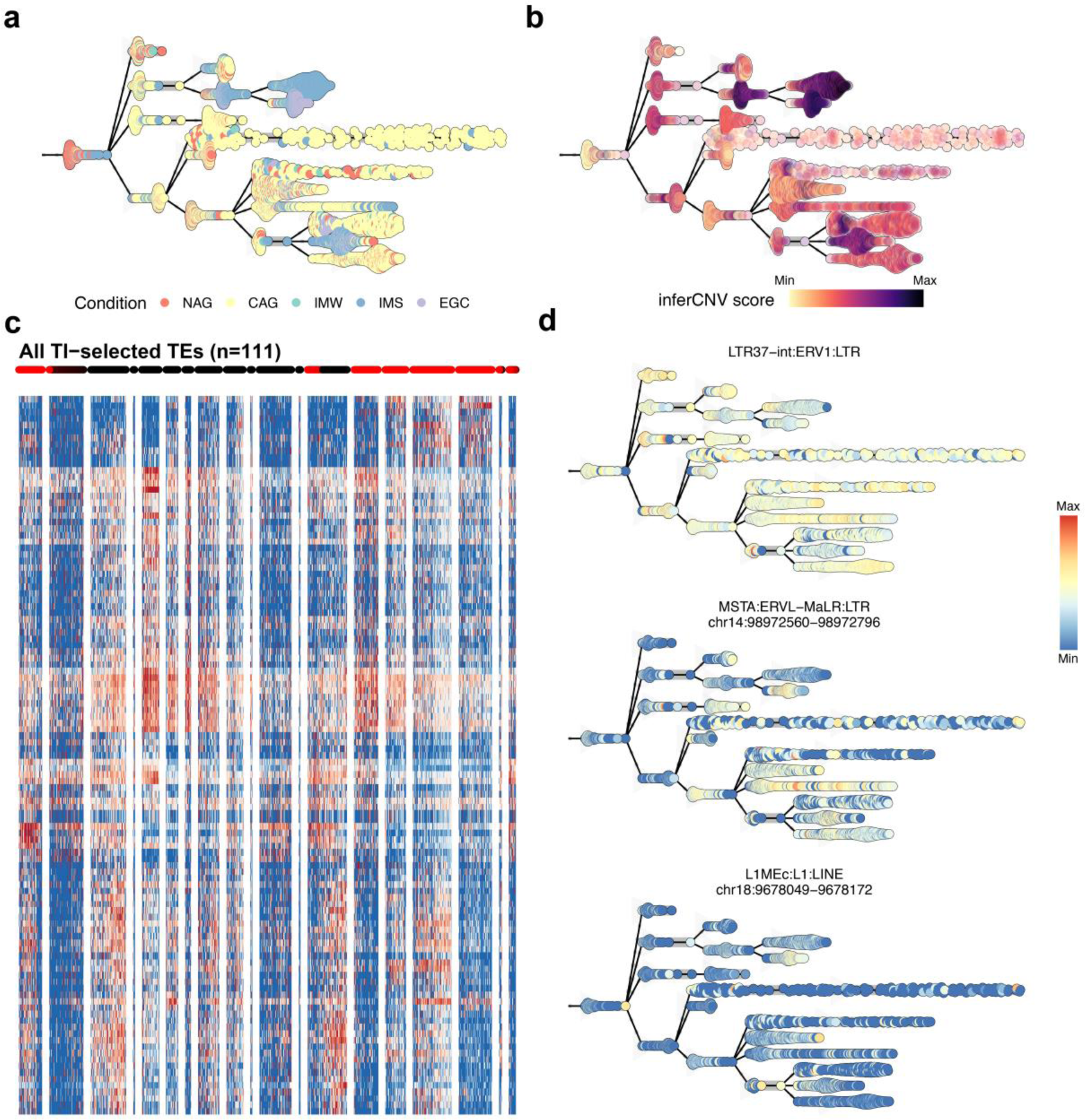
The single cell trajectory of EGC. **a.** Inferred pseudotemporal trajectory of the EGC dataset colored by condition: NAG in orange, CAG in yellow, IMW in cyan, IMS in light blue and EGC in purple. **b.** Pseudotemporal trajectory colored by inferCNV score: minimum values in light yellow, and maximum values in dark purple. **c.** Heatmap showing the expression of TEs associated with the increase in cell malignancy as measured by inferCNV scores. Colored dots above the heatmap correspond to a one-dimensional representation of the trajectory. Highlighted in red are the cells that go from the beginning of the trajectory to the Cancer milestone **d.** Example of TEs that are expressed in the malignant (i.e., high inferCNV score) branches of the trajectory.

The trajectory broadly recapitulates the progression from NAG to EGC: the majority of cancer cells appear in a lineage that originates from IMS cells, which in turn, originates from CAG cells. Notably, there are 2 milestones with a high number of IMS cells, with one of them continuing directly to the Cancer milestone. To add support to the trajectory in the context of cancer evolution, I also applied inferCNV ^41^ to generate per-cell scores of copy number variations (CNVs), which has been used as proxy of malignancy development ^13^. The inferCNV scores projected in the trajectory are also in concordance with the progression of CAG to IM, and subsequently to EGC (**Fig. 4b**). Cell type analysis of the trajectory shows that the two branches with the highest inferCNV scores depict the transitions between different cell types (**Supplementary Fig. 1**) The first branch reveals a transition from Enterocytes, to MSCs and Cancer, while the second branch reveals a transition from a milestone comprised by PMCs and some Enterocytes to one comprised by MSCs and Goblet cells. Although these cell types share a link on the basis of their transcriptional similarity ^6^, it is unclear whether they represent the actual cell evolution in EGC. In cancer, plasticity allows tumor cells to change between cell status ^42^. Thus, these observations might be indicative of cellular plasticity occurring during EGC, which would explain the cell type diversity in some branches with a malignant profile as revealed by their high CNV scores.

To better understand the impact of TE expression in the EGC cascade, the enrichment of TEs in the malignant milestones was assessed. This resulted in 111 TI-selected TEs (**Fig. 4c**), with 50 of them corresponding to markers of different cell populations and the remaining 61 without significant cell type specific expression (Supplementary data 3). Broadly speaking, three types of expression patterns throughout the trajectory were revealed by this analysis: 1. TE showing high and consistent expression (**Fig. 4d**, “LTR37-int”), 2. TE showing moderate levels of expression (**Fig. 4d**, “MSTA” located in chr14:98972560-98972796) and 3. TE expression mostly restricted to the malignant milestone (**Fig. 4d**, “L1MEc”, located in chr18:9678049-9678172). For example, “L1MEc” also appeared in the marker analysis as a Cancer cells-enriched TE (depicted in Fig. 3d), indicating some agreement between the 2 analyses. Notably, the 50 TEs that are also cell population markers all have locus resolution, while none of the other 61 have. The lack of locus resolution is usually indicative of expression of TE copies with a negligible number of discriminative mutations (i.e., evolutionary younger copies), hence the difficulty on accurately assigning reads to specific instances in the genome^25^. In this line, a possible scenario is that many of these copies become transcriptionally activated concurrently, pointing out to a pattern of global activation. For example, the widespread expression of “LTR37-in”, a TE belonging to the group of retroviruses, would be in agreement with evidence indicating that HERVs undergo global activation in cancer ^43^.

Collectively, this analysis showed that by leveraging TI methods, TEs are potentially contributing to the evolution of cancer cells and to the transition and interplay between cell status during EGC progression. In turn, the detection of TEs in the premalignant milestones might be informative to their role as biomarkers for the early detection of GC.

### The spatial expression of TEs in GC

Because I found evidence of TEs expressing from gastritis to EGC, I asked whether they are indeed expressed in GC. To this end, I analyzed spatially-resolved RNA-Seq data publicly available from a different study ^14^. The dataset corresponds to samples from 4 different patients, and was first analyzed with SoloTE and the resulting gene+TE expression matrices were processed with STutility ^44^. By taking advantage of the pathologist-annotated normal and tumor regions, the extent of TE expression occurring in the GC malignant regions was studied (**Fig. 5**).

**Fig. 5.**
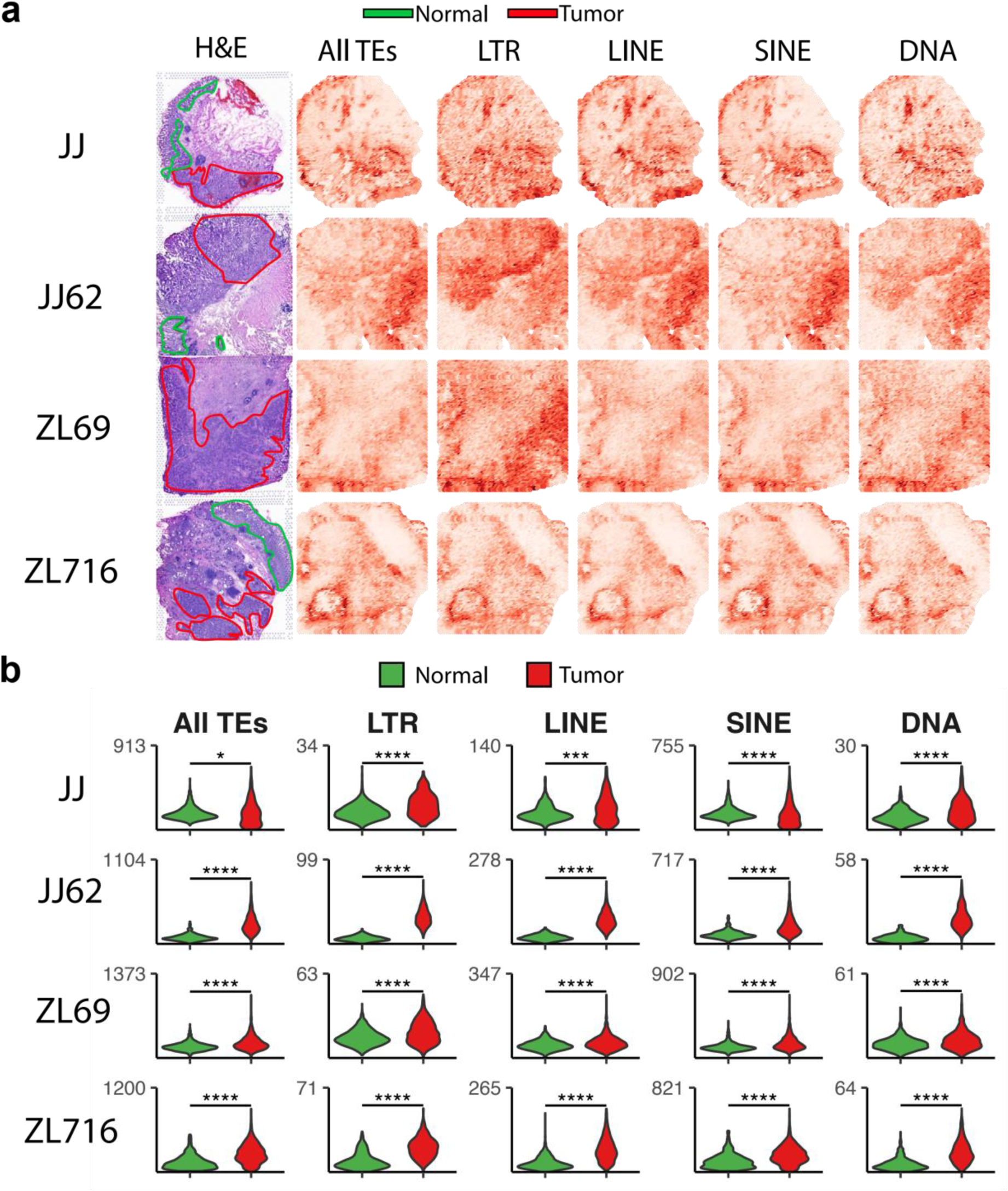
The spatial portrait of TE expression in GC. **a.** Spatially-resolved expression of TEs when analyzed collectively (“All TEs), or at the class level (LTR, LINE, SINE, DNA). “H&E” column shows the pathologist-annotated normal and tumor regions in green and red, respectively. **b.** Violin plots showing TE expression in the normal and tumor regions. Asterisks denote statistical significance at the following levels: **** - p ≤ 0.0001, *** - p>0.0001 and p ≤ 0.001, ** - p>0.001 and p ≤ 0.01, * - p > 0.01 and p ≤ 0.05.

Across all samples, global TE expression occurs in the tumor regions to varying degrees, and in all cases, the differences were statistically significant (**Fig. 5a** and **Fig. 5b**, “All TEs”). When observing the expression at the major TE type levels, subtle patterns can be seen. For example, a clear enrichment of LTRs in tumors is revealed (**Fig. 5**, “LTR”), while LINEs and SINEs exhibit higher expression in regions not annotated as normal nor as tumor (**Fig. 5**, “LINE” and “SINE”, respectively). Nonetheless, at the statistical level they are still relatively increased when comparing tumor versus normal regions (**Fig. 5b**). Interestingly, DNA TEs are also expressed, despite their lower presence in the human genome compared to retroelements. These results show that, similar to the single-cell part of this work, all major types of TEs become expressed in GC.

To investigate the spatial enrichment of specific TEs, I applied the FindMarkers function to compare expression in the different tissue regions using the normal epithelium as control. This analysis showed substantial variability in the number of spatially-enriched TEs detected on each sample: JJ62 – 148, JJ – 75, ZL716 – 57, ZL69 – 38 (**Supplementary Fig. 2**, Supplementary data 4). The number of TEs common between different combinations of samples was visualized in an upset plot (**Fig. 6a**). By using this result as guide, it was observed that 33 spatially-enriched TEs appear in at least 3 out of the 4 samples, and these were selected to build the set of “top” spatial TEs. Visualization of the TE type distribution indicates that despite the difference in number of enriched TEs detected on each sample, LINEs seem to be predominant, followed by SINEs, and then LTR and DNA (**Fig. 6b**). In the top set, DNA TEs do not appear, indicating that their activation follows the inter-patient GC heterogeneity captured in these samples. Conversely, there are 9 leading TEs in the top set because they were detected in all the samples (**Fig. 6c**). These 9 TEs correspond to 1 LTR, 4 LINEs and 4 SINEs, with 6 of them having locus resolution. In contrast with the single-cell analysis, these results show more sparsity in terms of the number of TEs whose location is detected unambiguously in the genome. For instance, in the total 33 TEs of the top set, only 15 (45.5%) have locus resolution, with 6 of them being amongst the leading 9 TEs present in all datasets. Nonetheless, when assessing the overlap between the top spatial TEs and the single-cell results, 29 (87.9%) out of the 33 top TEs were also detected in the single-cell analysis, with 22 (66.7%) of these TEs also associated with the malignant milestones detected in the trajectory analysis (**Supplementary Fig. 3**).

**Fig. 6.**
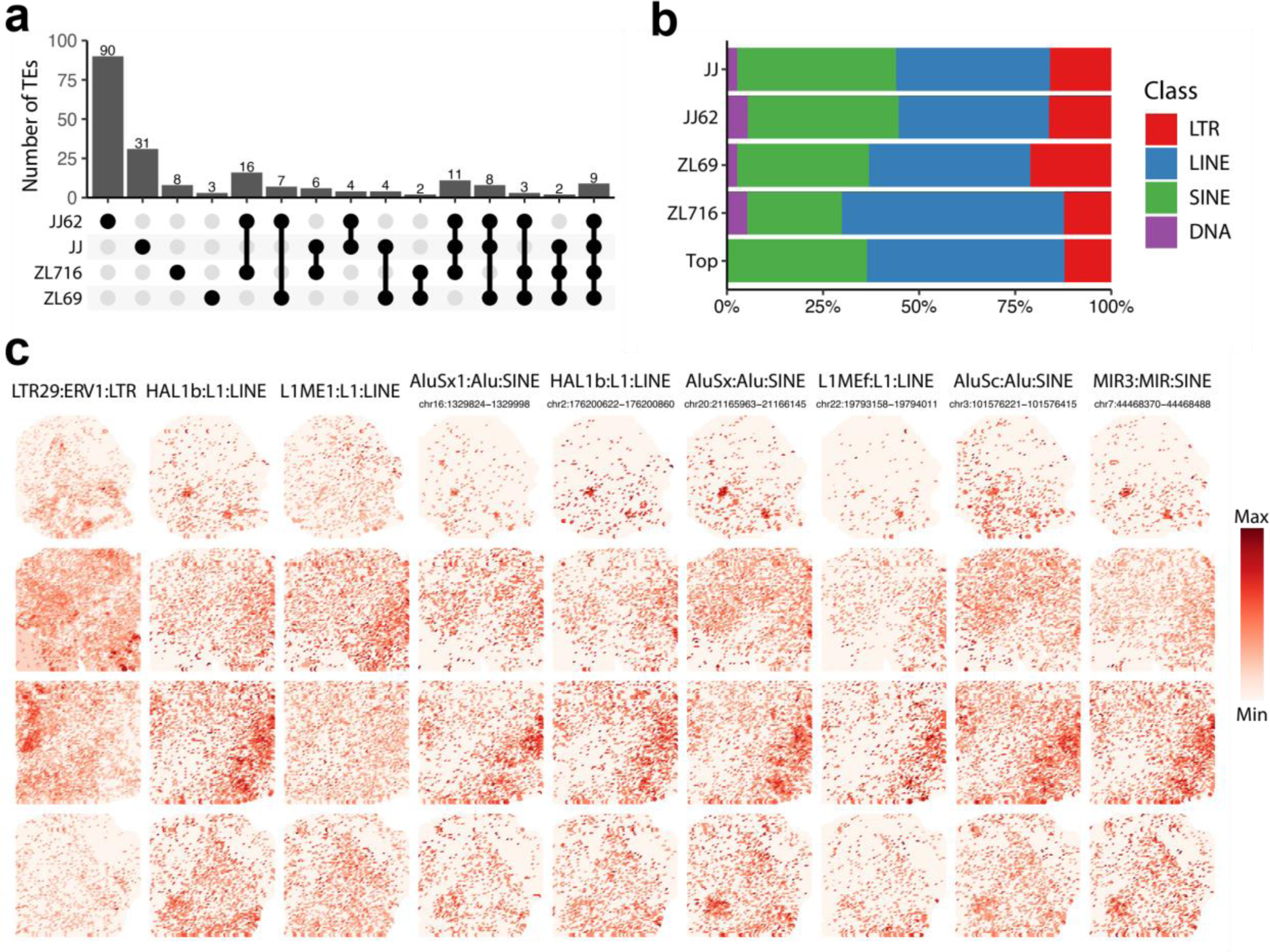
Spatially-enriched TEs. **a.** Upset plot of the spatially-enriched TEs. The number of TEs at each given overlap, represented by a set of connected dots in the lower half, is indicated as a bar plot. **b.** Class distribution of spatially-enriched TEs on each sample, and for the top TEs (“Top”) which corresponds to those identified in at least 3 out of the 4 samples. **c.** Feature plots of spatially-enriched TEs identified in all samples, depicting their expression across the tissue sections. For TEs identified with locus resolution, their genomic location is indicated under their name.

### The TE-mediated gene regulatory network during GC

To ask about the potential role of TEs as regulators of gene expression, a methodology similar to previous works ^13,43^ was applied: (1) using all the locus-specific TEs detected in both the single-cell and the spatial analysis, a gene-TE dictionary was built, considering all genes within 500 kbp of each TE; (2) Afterwards, to characterize the regulatory potential of TEs, the overlap with regulatory elements in GeneHancer and the ENCODE SCREEN candidate Cis-Regulatory Elements (cCREs) was assessed; (3) the gene-TE dictionary was filtered by considering either pre-defined interactions in GeneHancer or SCREEN, or if the gene was within 50 kbp of the TE; (4) finally, the Spearman correlation values were calculated for each gene-TE pair, and all interactions with correlation ≥ 0.3 were kept.

To get an overview of the genes potentially regulated by TEs, two additional analyses were carried out. First, an automated literature search using the NCBI E-Utilities ^45^ was carried out, and genes associated with publications in cancer were labeled as “Cancer gene”. Also, gene set enrichment analysis using the fgsea R package ^46^ was performed, using the Kyoto Encyclopedia of Genes and Genomes (KEGG) terms, and the Gene Ontology (GO) terms, and filtering results to those having adjusted p-value ≤ 0.05.

Following the aforementioned approach, a total of 3,151 interactions were predicted, with 1,142 of these classified as “TE GeneHancer”, 572 as “TE ENCODE cCREs”, and 1,437 as “TE coexpression” (Supplementary data 5). A total of 1,992 unique genes are regulated by TEs, with 573 of them labeled as “Cancer gene”. 210 unique genes (59 being Cancer genes) have 3 or more interactions with TEs, amounting to a total of 997 interactions: 445 “TE GeneHancer”, 96 “TE ENCODE cCREs” and 454 “TE coexpression” (**Fig. 7a**). Out of the 210 genes, 153 have either a TE GeneHancer or TE ENCODE cCREs interaction, providing further support to the hypothesis of TEs playing a regulatory role in GC. The remaining interactions met the correlation threshold, and the gene is in close genomic vicinity of the TEs, consistent with previous findings ^13,35^ suggesting such a regulatory role for TEs.

**Fig. 7.**
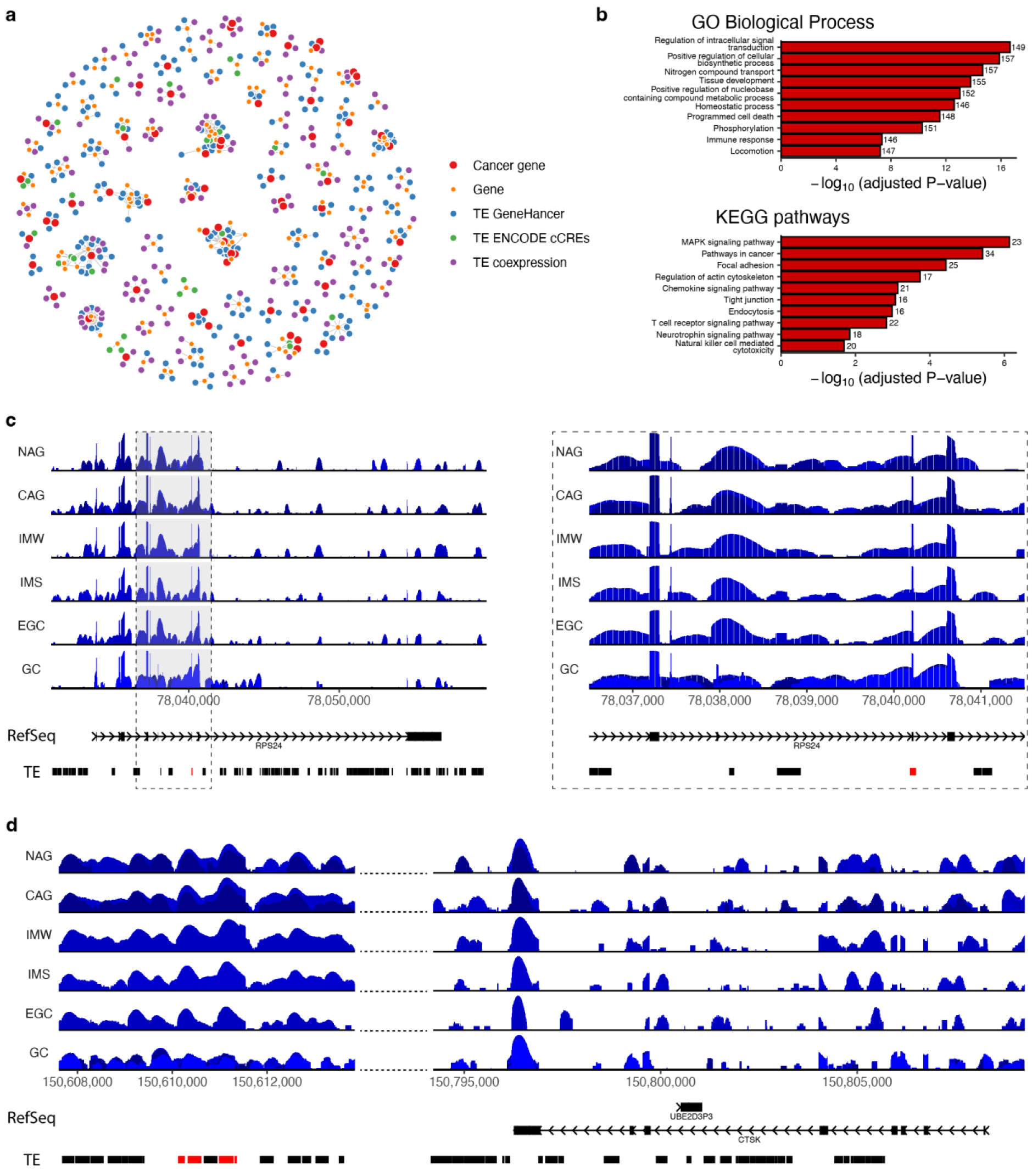
Network analysis of Gastric Cancer TEs. **a.** Regulatory network of genes associated with TEs. Genes previously associated with cancer are shown in red, and the remaining genes are shown in orange. TEs are colored according to their predicted regulatory link: TE GeneHancer in blue, TE ENCODE cCREs in green, and TE coexpression in purple. **b.** Top 10 enriched gene set terms for the Gene Ontology (GO) Biological Process category (upper half) and KEGG pathways category (lower half). **c.** Genome coverage plot of the intronic TE MamRTE1:RTE-BovB:LINE in locus chr10:78040182-78040254 (highlighted in red), having a predicted regulatory link with the RPS24 gene. On the right, the region enclosed with the dashed-lines rectangle is shown zoomed. **d.** Genome coverage plot of the region chr1:150610108-150809260, depicting upstream TEs (highlighted in red) having a predicted enhancer regulatory link with the CTSK gene.

At a wider scale, gene set enrichment analysis revealed many terms of relevance to cancer (**Fig. 7b**). For example, intracellular signal transduction, programmed cell death and phosphorylation are all processes long associated with cancer ^47–49^. Pathway-level analysis also depicts changes previously associated with cancer. The MAPK signaling pathway plays roles in cell proliferation, differentiation, migration and apoptosis, and is the top pathway likely regulated by TEs. Because of its role in many cell processes, malfunction on the pathway has been linked to cancer ^50^. “Pathways in cancer” is the second term, and it encompasses different pathways with evidence associating them with cancer. Thus, this result is in close agreement with the previous literature search analysis carried out independently. Another interesting example is “Endocytosis”, that has many functions in nutrient uptake. Disruptions in this pathway also play a significant role in cancer, and there is evidence pointing to a role in GC ^51,52^.

Finally, two examples of genes regulated by TEs are depicted: *RPS24* (**Fig. 7c**) and *CTSK* (**Fig. 7d**). *RPS24* was chosen because the interacting TE appears in all 3 analyses (single-cell, TI and spatial), has a partial overlap with one of its exons (**Fig. 7c**, TE highlighted in red), and the regulatory link is based on GeneHancer. In turn, the result would suggest that the TE is acting as an enhancer of the same gene. This would be similar to a documented event occurring in the *EGFR* gene, causing its up-regulation, which in turn, contributes to breast cancer ^53^. Also, over-expression of *RPS24* has been reported to be a biomarker of colorectal cancer ^54^. In these lines, the TE-mediated up-regulation proposed in this study might also highlight a role of *RPS24* in GC progression. On the other hand, *CTSK* has been previously implicated in GC, and the associated TEs are located about ∼200,000 kbp away from its locus, and have overlaps with ENCODE cCREs and GeneHancer elements, making this a good example of a long-distance regulatory link. *CTSK* has been reported as over-expressed in GC and has been proposed as GC biomarker ^55,56^. Additionally, it is unclear how it becomes over-expressed ^55^. This result would help bridge that gap, by indicating that up-regulation of *CTSK* is caused by TE-derived enhancers.

In sum, these results show that TEs are potentially regulating genes and pathways associated with cancer in several ways, strongly suggesting that TEs are implicated in the molecular aberrations that occur in GC.

## Discussion

GC is commonly detected at advanced stages, point in which the survival rate is low. When detected on initial stages, survival rate is high, prompting the need for better understanding the molecular mechanisms associated with its development and discovery of early biomarkers. In this work, I studied single-cell and spatial RNA data publicly available from different cohorts and provide evidence pointing to TEs as potential biomarkers and regulators of gene expression during GC.

Here, I showed that TE expression is a hallmark of the early GC cascade and during GC. The single-cell analysis revealed that thousands of TEs are cell-type enriched, and their expression increases as gastritis progresses to early GC. Although LTR ERVs have received more attention in cancer studies ^13,43^, there is increasing evidence depicting changes of several types of TEs in these malignances ^57^. Concordant with the latter evidence, in addition to LTRs, I also detected expression of LINE, SINE, and DNA TEs. Marker analysis revealed that TEs are enriched in the different cell populations to varying degrees. For example, the top Cancer markers also exhibit some level of expression in MSCs, in line with the proposed idea that MSCs might provide an environment for the origin of Cancer cells ^6^. Afterwards, the application of TI methods revealed a branching trajectory, broadly recapitulating the cascade from NAG, to CAG, to IM and to EGC. Particularly, some cell lineages show a clear association with the acquisition of the malignant status, based on their emergence in EGC and their high CNV score. Analysis of TE expression in the malignant lineages revealed 111 TEs that could potentially be early biomarkers for GC detection due to having expression not only in the later trajectory milestones, but in those leading up to them. Despite *dynverse* allowing a convenient assessment of many TI methods, the trajectory generated for this work still has some limitations, considering that in some instances it seemed to indicate cellular plasticity (phenomenon known to occur in cancer ^42^) rather than cell evolution. Nonetheless, the inferred trajectory is still informative and underlines an additional layer in which the study of TEs can provide insights to understanding GC progression.

Studying the spatial dataset generated from GC patients also confirmed TE expression. The importance of this is two-fold: first, it allowed bridge the gap between EGC and GC in terms of TEs, and second, it serves as additional and independent confirmation that changes in TE expression during GC development might indeed be representative of the malignancy. A caveat here is that differences were observed between patients, and overall, with the single-cell data. The diversity of TEs detected on each sample can be attributed to inter-patient and intra-tumor heterogeneity, and indeed in the original work it is reported that the regions with the most significant heterogeneity were selected for spatial sequencing ^14^. On the other hand, the lower TE detection observed in spatial data versus the single-cell data could be attributed to differences in the techniques: Visium spatial RNA-Seq captures about 5-10 cells per spot, and by sequencing a combination of cells could reduce the detection of several transcripts, including TE-derived ones. Nonetheless, almost 90% of the top spatial TEs were found in the single-cell data, and close to 70% are found in the list of TI-selected TEs, strongly suggesting that TE expression is a hallmark of GC, and that they might be playing a significant role in GC.

Finally, I adopted a gene-TE correlation approach to predict the impact of TEs in gene expression, which was coupled with the assessment of TEs as regulators via the overlap with GeneHancer and ENCODE cCREs. Close to 2,000 genes are likely regulated by TEs, and ∼500 of them have been previously linked with cancer. Gene enrichment analysis also depict the global impact of these potential regulatory events, and several of the top terms and pathways have been also associated with cancer. This gene-TE correlation approach is similar to the one applied in gallbladder cancer, where they validated the regulatory potential of selected TEs ^13^. There is a growing body of evidence supporting the role of TEs as regulators of gene expression either in their genomic vicinity or in long-distance locations by acting as enhancers ^23,29,43,57^, thus the findings reported here are in strong agreement with the idea of TEs involved in GC-related gene aberrations.

In conclusion, I show that TEs become activated during the progression of gastritis toward EGC, and in GC itself by leveraging datasets publicly provided by independent studies. I present evidence pointing that, in addition to becoming activated, TEs might influence gene regulation. In turn, this could contribute to the progression of GC. These findings highlight the biological and functional importance of studying TEs in this malignancy. The portrait of TE expression during GC development shown here advances our understanding of the disease, and pinpoints these elements as potential biomarkers for its early detection.

## Methods

### Raw sequencing data

The single-cell and spatial RNA-Seq data used in this study were obtained from publicly available databases. Single-cell FASTQ files were obtained from ^6^, made publicly available at the Sequence Read Archive (SRA) under accession SRP215370. On the other hand, Spatial data was obtained from ^14^, publicly available at the National Genomics Data Center Genome Sequence Archive (GSA) under accession HRA003070.

### TE expression analysis

To calculate TE expression on each dataset, the raw sequencing data was aligned to the human genome using STAR ^58^. First, the hg38 genome FASTA and ncbiRefSeq GTF annotation were downloaded from UCSC Genome Browser database ^59^ and used to generate the genome index. Then, the sequencing data of both the single-cell and spatial experiments was aligned with the following options to generate BAM files compliant with SoloTE (described later): *-- outSAMattributes NH HI nM AS CR UR CY UY CB UB GX GN sS sQ sM* to include cell barcode and UMI information in the output files, *--outFilterMultimapNmax 100 -- winAnchorMultimapNmax 100* to increase sensitivity of alignment to TEs, *--outSAMmultNmax 1 --outMultimapperOrder Random* to keep only one random alignment for multimapped reads, *--runThreadN 21* to set the number of process threads to 21 and *--runRNGseed 777* to set a random number generator seed to a fixed value for reproducibility. Each alignment file was then processed with SoloTE v1.09 ^28^, using the human genome hg38 version TE annotation in BED format obtained with the helper script *SoloTE_RepeatMasker_to_BED.py*. This process resulted in the raw count matrices that include TE expression.

### Single-cell analysis

Analysis of single-cell count matrices generated above was carried out using the Seurat v4.1.0 package ^60^ of the R statistical computing environment ^61^ version 4.1.1, as described next. First, single-cell matrices were filtered to keep only the quality-control filtered cells reported in the original work. Afterwards, the matrices were merged in a single object and processed using the default Seurat workflow. Briefly, the object was used as input to NormalizeData, FindVariableFeatures and ScaleData, in order to prepare it for principal component analysis using the RunPCA function. Then, 30 dimensions were used for FindNeighbors, and RunTSNE, and the clustering was obtained with FindClusters. Per-sample pseudobulk count matrices were generated with the AggregateExpression function, using the sample identifier as “group.id”. Then, the pseudobulked matrices were processed with DESeq2 ^32^, and the log-normalized counts were obtained with the rlog function, which were used to produce a per-sample PCA. For Figure 2, these steps were repeated 2 more times using a single-cell matrix with only genes and another one with only TEs.

To identify TEs whose expression increases in the progression towards early gastric cancer, DESeq2 was used again to test for differences between CAG, IM and EGC with respect to NAG, using adjusted P-value ≤ 0.05 as threshold for significance. In addition, to also include TEs highly expressed throughout all time points, those within the top 5% of expression were also selected. This list of EGC progression-associated TEs was used for marker analysis in the FindAllMarkers function to identify the specific cell populations in which they were enriched. Results of this step were also filtered using a threshold of adjusted P-value ≤ 0.05.

Trajectory Inference (TI) analysis was carried out using *dyno,* and related plots were generated using *dynplot*, both from the *dynverse* collection of R packages ^15^. The single-cell expression matrix was subsetted to the epithelial subtypes (PMC, GMC, Enteroendocrine, Neck-like, Chief, PC, Goblet, Enterocytes, MSCs, and Cancer) and then processed with the Seurat integration protocol. The “paga_tree” TI method was used for TI as it was identified as the most suitable for the dataset using the “guidelines_shiny” function. *inferCNV v1.8.1* ^41^ was run to generate per-cell copy number variation scores, which were used as an additional validation of the inferred trajectory. The “branch_feature_importance” function was used to assess TEs enriched in the cell lineages associated with cancer progression.

### Spatial analysis

Spatial data was processed with STutility v1.1.1 ^44^, which is built in top of the Seurat package. First, for each of the 4 samples an object was created, and processed with the SCTransform function to obtain the normalized expression. Then, total TE and per-class (LTR, LINE, SINE and DNA) TE expression was calculated by aggregating all the normalized counts respectively. Statistical differences in TE expression between tumor and normal regions across the tissue were tested using the Wilcoxon test implemented in the base R function *wilcox.test*. The results of this step were depicted in the violin plots of Figure 5b, highlighting the Wilcoxon test p-values obtained.

To find TEs with higher expression in tumor and non-normal regions of the tissues, differential expression analysis was carried out with the FindMarkers function. This was done by taking advantage of the pathologist annotations, using the normal epithelium regions as control (or the unannotated region in the case of sample ZL69) and each of the remaining regions as test groups. All TEs with adjusted p-value ≤ 0.05 were then selected as spatially-enriched. The overlap between the spatially-enriched TEs detected in each sample was assessed via an upset plot generated with the ggupset v0.3.0 package ^62^.

### Network analysis

To build the TE-gene networks, a list of TEs was built from those enriched in the single-cell or spatial data. Selected single-cell TEs were those associated with the progression from NAG to EGC and that also appear as markers of cell populations, whereas selected spatial TEs were those found with spatial enrichment in at least 3 out of the 4 samples.

The selected TEs were then processed to identify potential regulatory links on the basis of their genomic location, using a methodology similar as previously published works ^13,43^. First, interactions between regulatory elements and genes were obtained from GeneHancer and the SCREEN database. Particularly, GeneHancer v4.7 interactions were downloaded from https://genecards.weizmann.ac.il/geneloc/index.shtml, and the ENCODE SCREEN Registry of candidate Cis-Regulatory Elements (cCREs) V3 ^63^ from https://screen.encodeproject.org/. Afterwards, a TE-gene dictionary was built containing all genes within 500 kbp from the selected TEs. Using BEDTools v2.30.0 ^64^, the overlap between selected TEs and regulatory elements was assessed to filter and classify the TE-gene dictionary into potential interactions: “TE GeneHancer”, if the TE overlapped with GeneHancer elements and the gene was a target of the regulatory element, “TE ENCODE cCREs” if the TE overlapped didn’t overlap with GeneHancer elements, but overlapped with ENCODE cCREs and the gene was a target of the regulatory element, and finally into “TE coexpression” if they didn’t have any overlap with regulatory elements, but the TE and gene were within 50 kbp of each other. Finally, the Spearman correlation between each TE-gene pair in the dictionary was calculated and those pairs with correlation ≥ 0.3 were selected to build the TE-gene network depicted in Figure 7.

### Gene set enrichment analysis

Gene set enrichment analysis was carried out as described before ^43^, using the “geseca” function of the fgsea v1.29.1 R package ^46^. The genes used as input were those selected at the final step of the network analysis, and the following parameters were specified: *minSize = 1, maxSize = 500, nPermSimple=10000* and *center=FALSE, scale=FALSE*. The analysis was carried out twice, first using category “C2”, which correspond to Kyoto Encyclopedia of Genes and Genomes (KEGG) terms, and then using “C5”, which corresponds to Gene Ontology (GO) terms. Enriched terms having adjusted p-value ≤ 0.05 were selected.

### Plots

*ggplot2* v3.4.2 ^65^ was used to generate the Figure 2a PCA plots, heatmap in Figure 3a, bar plots in Figure 3b, Figure 6b, and Figure 7b, and violin plots in Figure 5b. The *ggcoverage* v1.2.0 R package ^66^ was used to plot the RNA-Seq coverage at regions chr10:78032363-78058306 and chr1:150610108-150809260 shown in Figure 7c and 7d. The region chr10:78032363-78058306 depicts the *RPS24* gene and an intronic TE, whereas the region chr1:150610108-150809260 depicts a potential long-range interaction between an upstream TE and the *CTSK* gene.

## Data availability

The single-cell data is publicly available at Sequence Read Archive (SRA) under accession SRP215370. The spatial data is publicly available at the National Genomics Data Center Genome Sequence Archive (GSA) under accession HRA003070.

## Author contributions

B.V.M. conceived the project. B.V.M. initiated the project and wrote the paper. B.V.M performed the bioinformatic analysis. B.V.M. supervised and funded the project.

